# Dietary cholesterol reduces blood pressure and alters lipid profiles in stroke-prone spontaneously hypertensive rats

**DOI:** 10.64898/2026.01.27.702179

**Authors:** Yutaro Nishikata, Kenjiro Tatematsu, Yoshiaki Saito, Toshiyuki Matsunaga, Naoki Ohara

## Abstract

Although cholesterol (Chol) is widely recognized as a risk factor for cardiovascular disease, dietary Chol intake has been reported to extend the lifespan of stroke-prone spontaneously hypertensive rats (SHRSP). The mechanisms responsible for this paradoxical effect remain unclear. The present study examined changes in organ lipid profiles and associated molecular factors in SHRSP rats fed a Chol-enriched diet. Four-week-old male SHRSP/Izm rats were assigned to three groups and fed ad libitum for 12 weeks with either a control diet (Ctr), a diet supplemented with 1% w/w Chol (Chol), or a diet containing 1% w/w Chol plus 0.025% w/w lovastatin (Stt) to suppress endogenous Chol synthesis. Systolic blood pressure was measured before and after the feeding period, and tissues were collected for analyses of sterol content, fatty acid composition, prostaglandin E₂ (PGE_2_) levels, and renal histopathology.

Relative to the Ctr group, the Chol group exhibited a significant 9–10% reduction in systolic blood pressure. This reduction was accompanied by pronounced alterations in lipid profiles, including changes in phytosterol content and decreased arachidonic acid ratios in serum and kidney. There was a downward trend in hepatic PGE_2_ levels, and a similar tendency was observed in the kidney. Comparable changes in lipid profiles were observed in the Stt group. Histological analysis revealed modest attenuation of renal pathological features in Chol-fed rats.

This study demonstrates for the first time that dietary Chol reduces renal phytosterol accumulation and suppresses the AA-PGE₂ axis, changes that coincide with a 9–10% reduction in systolic blood pressure and attenuated glomerular inflammation. These integrated findings provide a mechanistic framework linking dietary Chol to the previously reported lifespan extension in this stroke-prone model. Although these changes may contribute to improved renal pathology, further studies are required to clarify causal relationships.

## Introduction

Sterols and fatty acids (FAs) are prevalent dietary lipids that serve essential physiological functions. Cholesterol (Chol), which is predominantly obtained from animal fats and oils, constitutes a fundamental component of animal cell membranes and functions as a precursor for steroid hormones and bile acids [1]. Elevated blood Chol levels are widely acknowledged as a significant risk factor for atherosclerotic cardiovascular diseases, primarily due to the propensity for the formation of Chol-rich plaques [2]. In contrast, epidemiological studies and meta-analyses have consistently demonstrated an inverse correlation between serum Chol levels and the risk of hemorrhagic stroke [3]. The mechanisms underlying this paradoxical association remain insufficiently characterized.

Stroke-prone spontaneously hypertensive rats (SHRSP) have been established as a model for human hypertension and hemorrhagic stroke [4]. In this strain, the lifespan is curtailed by fatal strokes that result from severe hypertension associated with renal impairment [5]. Notably, SHRSP rats exhibit a paradoxical response to dietary lipids; diets rich in plant-derived phytosterols (PS) shorten the survival by increasing blood pressure [6–8], whereas a Chol-rich diet prolongs lifespan [26]. This strain is characterized by a mutation in the ATP-binding cassette transporter G5 (ABCG5), resulting in accelerated PS accumulation within target tissues [9]. The accumulated PS has been shown to partially replace membrane Chol, thereby reducing membrane integrity and increasing vascular tissue fragility [10]. Under the severe hypertension characteristic of SHRSP rats, these fragile membranes, particularly in the brain, are prone to rupture, resulting in hemorrhagic stroke [11].

Chol and PS have been found to share similar intestinal absorption pathways. Both sterols have been observed to form micelles with bile acids and are taken up into epithelial cells via Niemann-Pick C1-like 1 (NPC1L1) [12]. After this process, approximately 50% of Chol and 90% of PS are re-excreted into the small intestinal lumen by the ATP-binding cassette transporter G5/8 (ABCG5/8) [13]. Consequently, PS exerts a competitive inhibition on Chol absorption, thereby demonstrating a Chol-lowering effect in the blood. Conversely, statins (Stt), which function as inhibitors of 3-hydroxy-3-methylglutaryl-CoA (HMG-CoA) reductase (HMGCR), represent the most prevalent treatment modality for hypercholesterolemia in clinical practice [14]. In addition to its capacity to reduce Chol, Stt exerts several lipid-lowering–independent effects. These include the stabilization and attenuation of oxidative stress, vascular inflammation, and blood pressure elevation [15–17]. Although Stt have been reported to improve survival and reduce stroke-related outcomes in SHRSP rats [18], their effects under conditions of excessive dietary Chol intake have not been fully examined.

Chol and FA metabolisms are closely interconnected, and dietary Chol influences lipid metabolism through transcriptional regulators such as sterol regulatory element-binding proteins (SREBPs) [19]. Polyunsaturated FAs (PUFAs), specifically those belonging to the n-6 and n-3 series, have been identified as playing pivotal roles in the regulation of inflammation [20]. Specifically, arachidonic acid (AA) (20:4n-6) is metabolized into prostaglandins (PGs) and other compounds that regulate inflammatory responses [21]. Therefore, alterations in sterol composition may exert an influence on FA-derived inflammatory pathways. However, the mechanisms by which dietary Chol intake modifies sterol composition and associated FA metabolic pathways in SHRSP rats remain to be elucidated, as do the relationships between these changes and the pathological features characteristic of this strain. Chol and PS are ingested daily, and Stt is frequently utilized in clinical settings. Consequently, elucidating the underlying mechanisms through which these interactions induce pathologies in cerebrovascular disease models offers significant insights from both nutritional and clinical perspectives.

The objective of this study was to elucidate the impact of dietary Chol and Stt on sterol and FA metabolism in SHRSP rats, and to determine the role of these metabolic changes in the pathological characteristics that are hallmarks of this strain.

## Materials and Methods

### Test diets

5001 Laboratory Rodent Diet Meal (PMI Nutrition International, Arden Hills, MN) was procured as the basal diet. The experiment involved the preparation of three test diets: a control (Ctr) group, a Chol group, and a Stt group. The experimental diets for the Chol and Stt groups were prepared by mixing the basal diet with 1% w/w Chol (FUJIFILM Wako Pure Chemical Co., Osaka, Japan) or with 1% w/w Chol and 0.025% w/w lovastatin (Tokyo Chemical Industry Co., Tokyo, Japan), respectively. The diets were stored at temperatures below -20°C until use. Tables 1 and 2 present the compositions of fatty FA and sterol contents of the diets.

**Table 1.** FA compositions and total levels of the test diets.

| **FA(%)** | **Ctr** | **Chol** | **Stt** |
| --- | --- | --- | --- |
| **14:0** | 1.5 | 1.5 | 1.5 |
| **16:0** | 21.9 | 22.0 | 21.9 |
| **16:1** | 2.1 | 2.3 | 2.2 |
| **18:0** | 7.6 | 7.5 | 7.7 |
| **18:1n-9** | 30.4 | 30.5 | 30.2 |
| **18:1n-7** | 1.5 | 1.4 | 1.6 |
| **18:2n-6** | 30.4 | 30.1 | 30.1 |
| **18:3n-6** | 0.1 | 0.1 | 0.1 |
| **18:3n-3** | 2.4 | 2.3 | 2.4 |
| **20:0** | 0.5 | 0.5 | 0.5 |
| **20:1n-9** | 0.1 | 0.1 | 0.1 |
| **20:5n-3** | 0.8 | 0.8 | 0.9 |
| **22:0** | 0.1 | 0.1 | 0.1 |
| **22:6n-3** | 0.6 | 0.7 | 0.8 |
| **24:0** | 0.1 | 0.1 | 0.1 |
| **Total FA (mg/g diet)** | 44.5 | 44.5 | 43.0 |

Fatty acids (FA) in each diet were derived from the control diet (5001 Laboratory Rodent Diet Meal). Values are presented as means (n = 2). Abbreviations: Ctr, control group; Chol, cholesterol group; Stt, statin group.

**Table 2.** Sterol levels of the test diets.

| **Sterol (mg/g diet)** | **Ctr** | **Chol** | **Stt** |
| --- | --- | --- | --- |
| **Cholesterol** | 0.293 | 10.599 | 10.394 |
| **Campesterol** | 0.075 | 0.081 | 0.083 |
| **Stigmasterol** | 0.028 | 0.031 | 0.032 |
| **β-Sitosterol** | 0.285 | 0.283 | 0.289 |
| **Total phytosterol** | 0.388 | 0.395 | 0.404 |

The Chol and Stt diets were supplemented with 1% w/w Chol. All phytosterols in each diet were derived from the control diet (5001 Laboratory Rodent Diet Meal). Values are presented as means (n = 2). Abbreviations: Ctr, control group; Chol, cholesterol group; Stt, statin group.

### Animals

The present study was conducted in strict accordance with the recommendations set forth in the Guide for the Care and Use of Laboratory Animals of the National Institutes of Health [22]. The study protocol was approved by the Institutional Animal Care and Use Committee of Gifu Pharmaceutical University (Gifu, Japan; Permit No. 2023-058) and the Committee for Animal Research and Welfare of Gifu University (Permit No. 2023-046). Terminal procedures were performed under deep anesthesia, followed by euthanasia, and all efforts were made to minimize animal suffering. All rats were maintained under specific pathogen-free conditions at 23°C ± 3°C and 55% ± 15% humidity in a 12:12-h light/dark cycle.

Eighteen male SHRSP/Izm rats, aged four weeks, were procured from Japan SLC Inc. (Shizuoka, Japan). Following acclimatization, the animals were randomly assigned to three groups (Ctr, Chol, and Stt; n = 6 per group) and housed in groups of three per cage. Animals in each group were permitted unrestricted access to the test diet and drinking water for a period of 12 weeks. Body weight and food consumption were monitored on a weekly basis. The subjects’ food intake was meticulously monitored using a specialized feeding device. Furthermore, blood pressure was measured before and after the feeding period using a tail-cuff method on a blood pressure monitor (MK-2000ST, Muromachi Kikai Co., Tokyo, Japan). At the conclusion of the feeding period, the animals were anesthetized by inhalation of isoflurane. Following euthanasia, the major organs were promptly harvested, weighed, frozen in liquid nitrogen, and stored at -80°C until use.

### Lipid analysis

The composition of the test diet, serum, liver, and kidney in terms of FA and sterol was determined according to the method described in our previous report, with certain modifications [7]. In brief, total lipids were extracted using the Bligh and Dyer method [23], and FAs were converted to methyl esters using a 10% HCl-methanol solution. (Tokyo Chemical Industry Co.). Following the extraction of methyl esters by means of petroleum ether, gas chromatography (GC-2010; Shimadzu Co., Kyoto, Japan), equipped with a DB-225 capillary column (Agilent Technologies, Inc., Santa Clara, CA), for the analysis of FAs. Heptadecanoic acid was utilized as the internal standard. The composition of free FA in the kidney was measured after the separation of phospholipids (PL) and neutral lipids, including free FA, by solid-phase extraction using the NH2 column (Bond Elut NH2; Agilent Technologies, Inc.).

For the purpose of sterol analysis, each tissue specimen was subjected to an incubation process with a 20% potassium hydroxide ethanol solution at a temperature of 100°C for a duration of two hours. This process, known as saponification, involved the use of betulin as an internal standard. The sterol fraction was extracted with hexane and converted to a trimethylsilyl derivative using TMS-HT^®^ reagent (Tokyo Chemical Industry Co.). The quantification of these levels was performed by gas chromatography (GC) using a capillary column (DB-1; Agilent Technologies, Inc.).

### Quantitative analysis of mRNA expression

Total RNA was extracted from liver tissue using TRIzol™ reagent (Thermo Fisher Scientific Inc., Madison, WI) according to the manufacturer’s instructions. RNA samples from each group (n = 6) were then pooled and treated with DNase I (Promega Corporation, Madison, WI). Real-time RT-PCR was performed using the RNA-direct^®^ SYBR™ Green Realtime PCR Master Mix (Toyobo Co., Osaka, Japan) on a StepOne Real-Time PCR System (Thermo Fisher Scientific Inc.). Each pooled RNA sample was analyzed in duplicate, and the experiment was independently repeated three times. Gene expression was calculated using the ΔΔCt method and normalized to β-actin. The primer sequences utilized for RT-PCR are enumerated in Table 3.

**Table 3.**
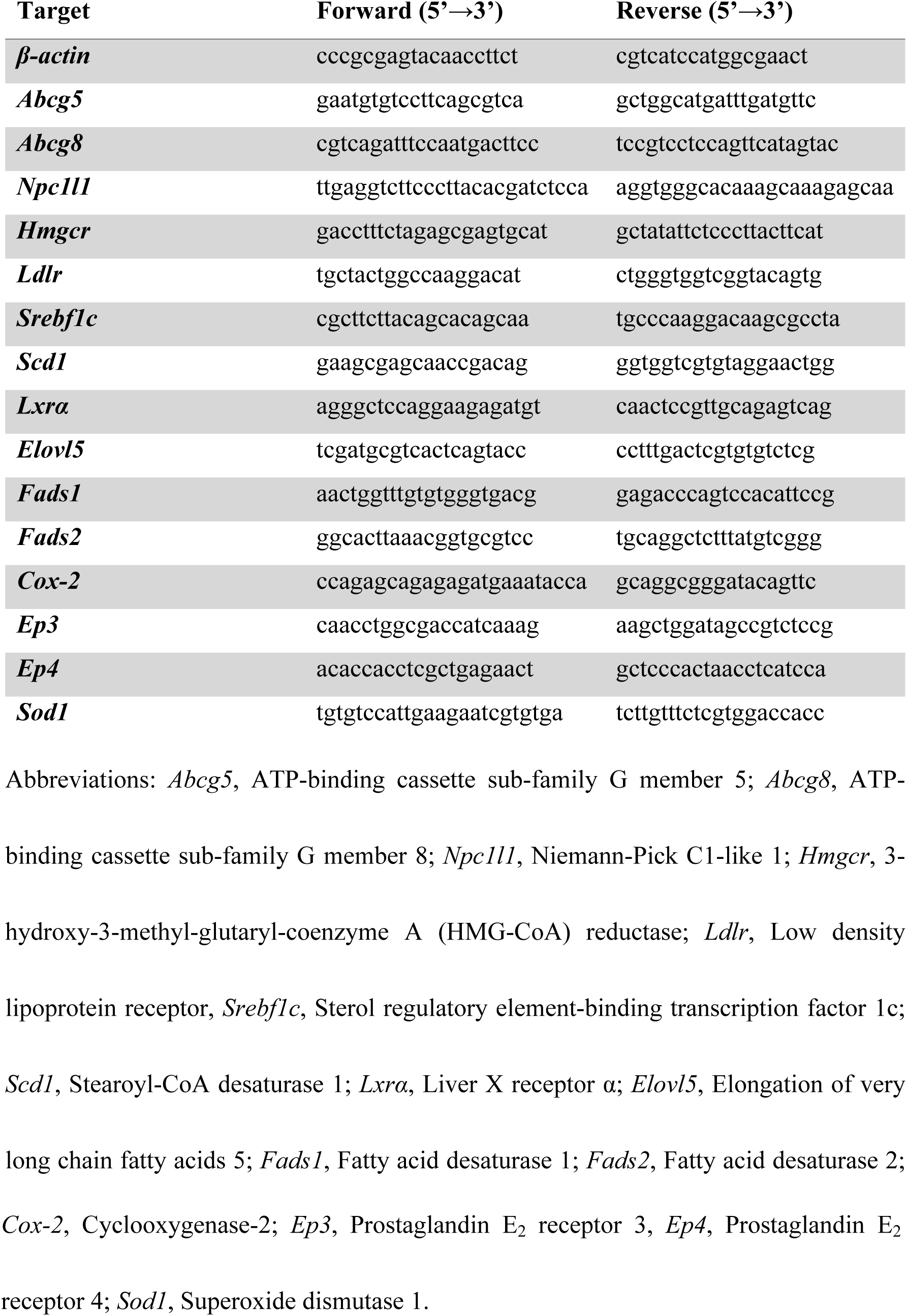
The primer sequences used for RT-PCR.

### Biochemical assay

The concentrations of prostaglandin E_2_ (PGE_2_) in the liver and kidneys were evaluated using the PGE_2_ Monoclonal ELISA Kit (Cayman Chemical Co., Ann Arbor, MI).

The activity of superoxide dismutase (SOD) in serum, liver, and kidney was measured using the SOD Assay Kit - WST (Dojin Molecular Technology Co., Kumamoto Prefecture) after pooling each tissue. All procedures were performed in strict accordance with the instructions provided with the kits.

### Histological analysis

Kidney tissue samples were fixed using 10% formalin neutral buffer solution and embedded in paraffin. The samples were then cut into 4-μm horizontal sections and stained with hematoxylin and eosin for histological examination. The process of embedding and staining was subcontracted to Morphotechnology Co. (Hokkaido, Japan).

A pathologist then compared each sample with normal tissue and assigned scores using a five-grade scoring system: – (no abnormal change), ± (very slight change), + (slight change), ⧺ (moderate change), and ⧻ (marked change).

### Statistical analysis

Statistical analysis was performed using KyPlot v.6.0 software (Keyence, Osaka, Japan). All data are expressed as the mean ± standard deviation (SD). A one-way analysis of variance (ANOVA) was used to compare the three feed groups, except for food intake and SOD activity. This was followed by a Tukey’s multiple comparison test as a post hoc test. In all cases, the significance level was set at p < 0.05.

## Results

### Body weight, food intake, blood pressure, and organ weight

No significant differences in body weight and food intake were observed among the three groups throughout the 12-week feeding period (Fig. 1A and B). A significant decrease in systolic blood pressure (SBP) was observed in the Chol (223 ± 6 mmHg) and Stt (220 ± 11 mmHg) groups compared with the Ctr (240 ± 11 mmHg) group following the 12-week feeding period (Fig. 2). Table 4 presents the organ weight at the time of sacrifice. Liver weight was significantly higher in the Chol and Stt groups than in the Ctr group. However, no significant differences were observed in the weights of the other organs.

**Fig 1.**
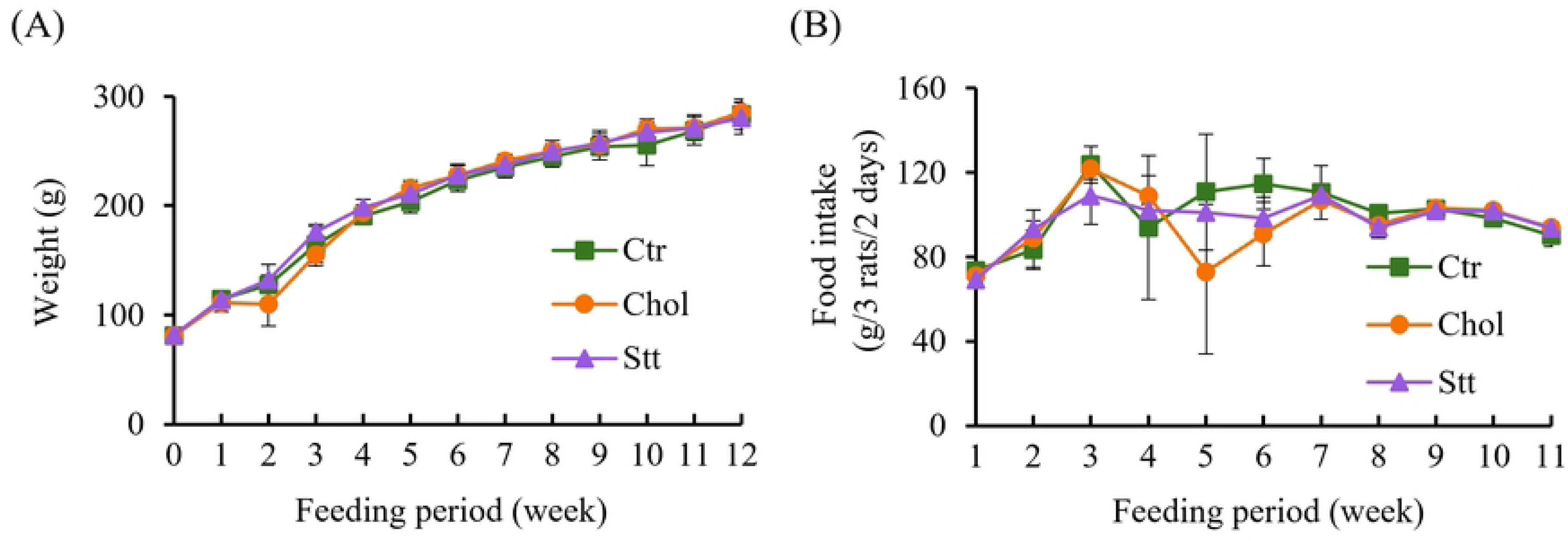
Body weight and food intake of the SHRSP rats. (A) Body weight and (B) food intake during the experimental period. Values are presented as mean ± SD (n = 6/group for body weight; n = 2 cages/group for food intake). No significant differences were observed among the three groups. Abbreviations: Ctr, control group; Chol, cholesterol group; Stt, statin group; SD, standard deviation; SHRSP, stroke-prone spontaneously hypertensive rats.

**Fig 2.**
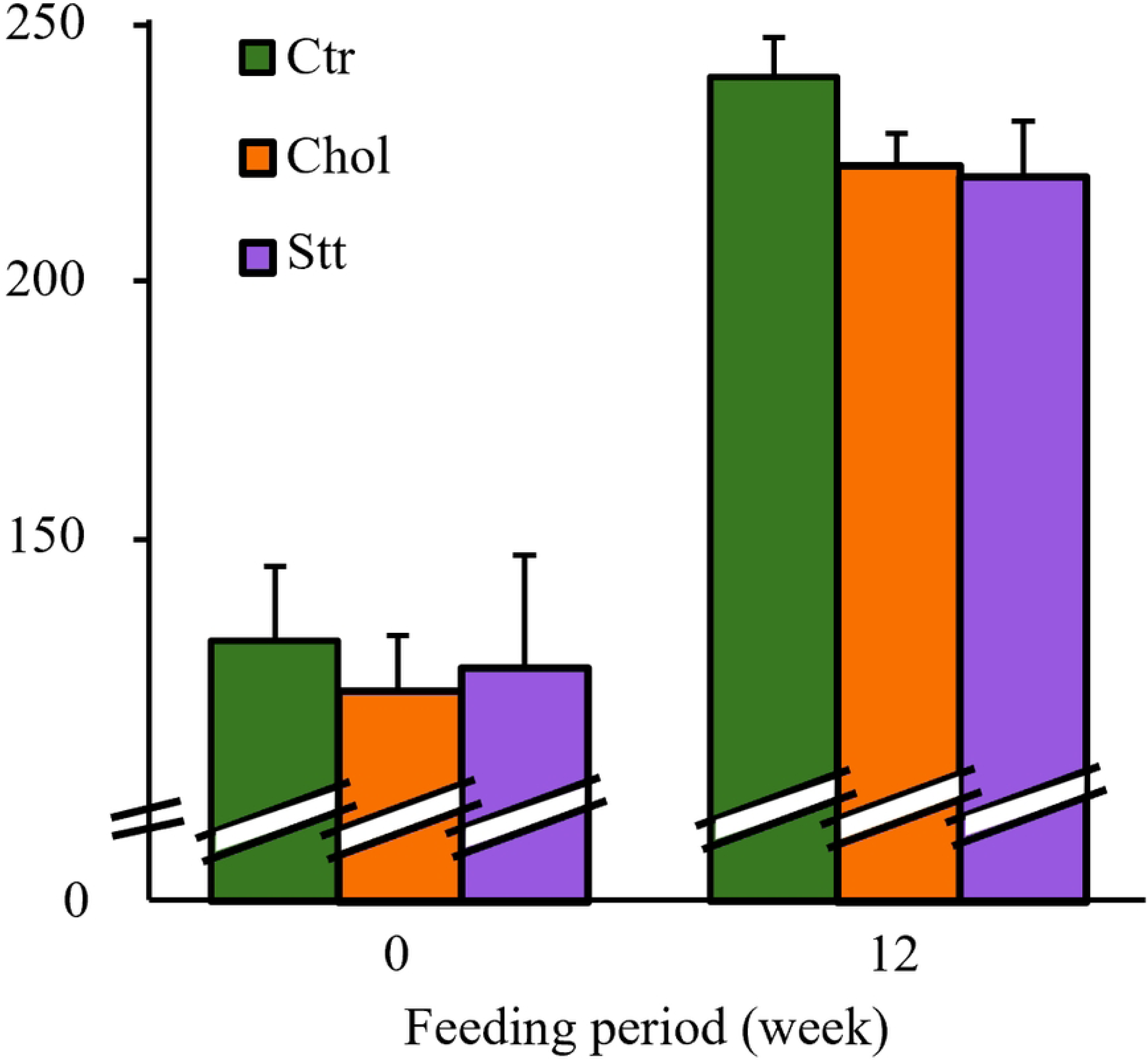
Systolic blood pressure of the SHRSP rats. Values are presented as mean ± SD (*n* = 6/group). \**p* < 0.05 vs. Ctr. Abbreviations: Ctr, control group; Chol, cholesterol group; Stt, statin group; SD, standard deviation; SHRSP, stroke-prone spontaneously hypertensive rats.

**Table 4.** Organ weights of SHRSP rats.

| Weight (g) | Ctr | Chol | Stt |
| --- | --- | --- | --- |
| Liver | 9.07 ± 0.83 | 11.27 ± 0.66* | 11.35 ± 1.20* |
| Kidney (Left) | 1.11 ± 0.08 | 1.09 ± 0.06 | 1.12 ± 0.07 |
| Kidney (Right) | 1.07 ± 0.11 | 1.09 ± 0.07 | 1.09 ± 0.06 |

Values are presented as mean ± SD (n = 6/group). \**p* < 0.05 vs. Ctr. Abbreviations: Ctr, control group; Chol, cholesterol group; Stt, statin group; SD, standard deviation; SHRSP, stroke-prone spontaneously hypertensive rats.

### Lipid analysis

An examination of sterol levels and FA compositions in the liver, serum, and kidney was conducted to ascertain the effects of dietary Chol and Stt in SHRSP rats. The consumption of Chol resulted in elevated levels of Chol in the liver, as compared with the control diet. Conversely, serum Chol levels were reduced in the Stt group relative to the Chol group (see Figures 3A and 3B). Renal Chol levels did not significantly differ among the three diet groups (Fig. 3C). In relation to PS, resulted in an increase in hepatic campesterol (Camp) and total PS levels, while decreasing total PS levels in both serum and kidney (Fig. 3A-C). A comparative analysis revealed a decrease in β-sitosterol (Sito) levels across all the tissues examined within the Chol group when contrasted with the Ctr group. Conversely, Stt supplementation led to a decrease in hepatic Camp and total PS levels relative to the Chol diet, while kidney sterol levels remained largely unaltered.

**Fig 3.**
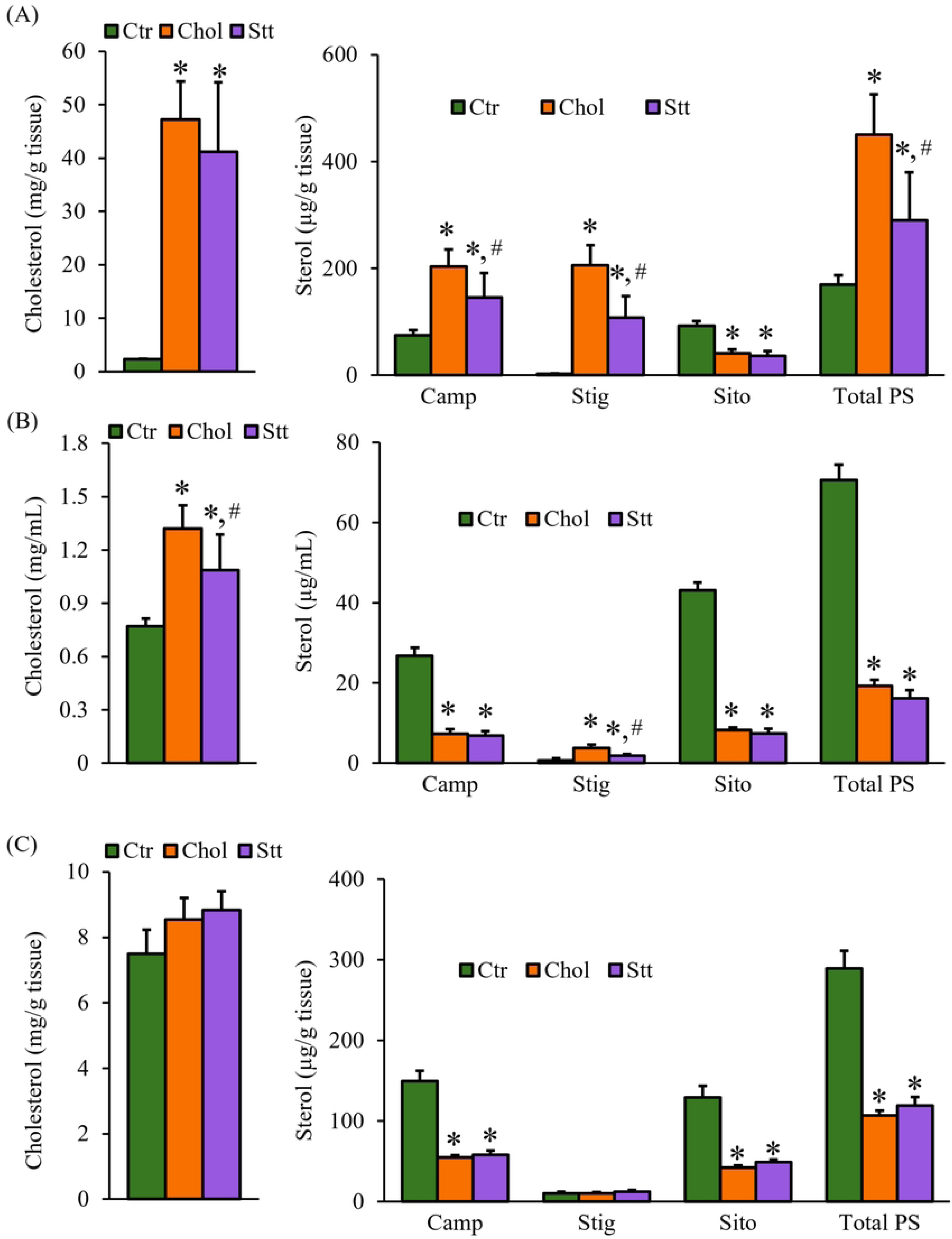
Sterol levels in the liver, serum, and kidney of SHRSP rats. (A) Hepatic, (B) serum and (C) renal Chol levels (left row) and PS levels (right row). Values are presented as mean ± SD (*n* = 6/group). \**p* < 0.05 vs. Ctr; ^#^*p* < 0.05 vs. Chol. Abbreviations: Ctr, control group; Chol, cholesterol group; Stt, statin group; Camp, campesterol; Stig, stigmasterol; Sito, β-Sitosterol; PS, phytosterol; SD, standard deviation; SHRSP, stroke-prone spontaneously hypertensive rats.

As illustrated in Table 5, there was a significant increase in the total FA level in the liver among the Chol and Stt groups in comparison with the Ctr group. With respect to the composition of FA, an increase in palmitoleic acid (16:1n-7), oleic acid (18:1n-9), and linoleic acid (18:2n-6), accompanied by a decrease in palmitic acid (16:0), stearic acid (18:0), and AA, was observed in both the liver and serum of the Chol group in comparison with the Ctr group (Tables 5 and 6). In the kidney PL, there was an increase in 18:2n-6 and a decrease in AA (see Table 7). In the Stt group, AA composition exhibited a decrease in the serum, liver, and kidney PL compared with the Ctr group. Notably, among n-6 PUFAs, a consistent reduction in AA was observed across multiple tissues.

**Table 5.**
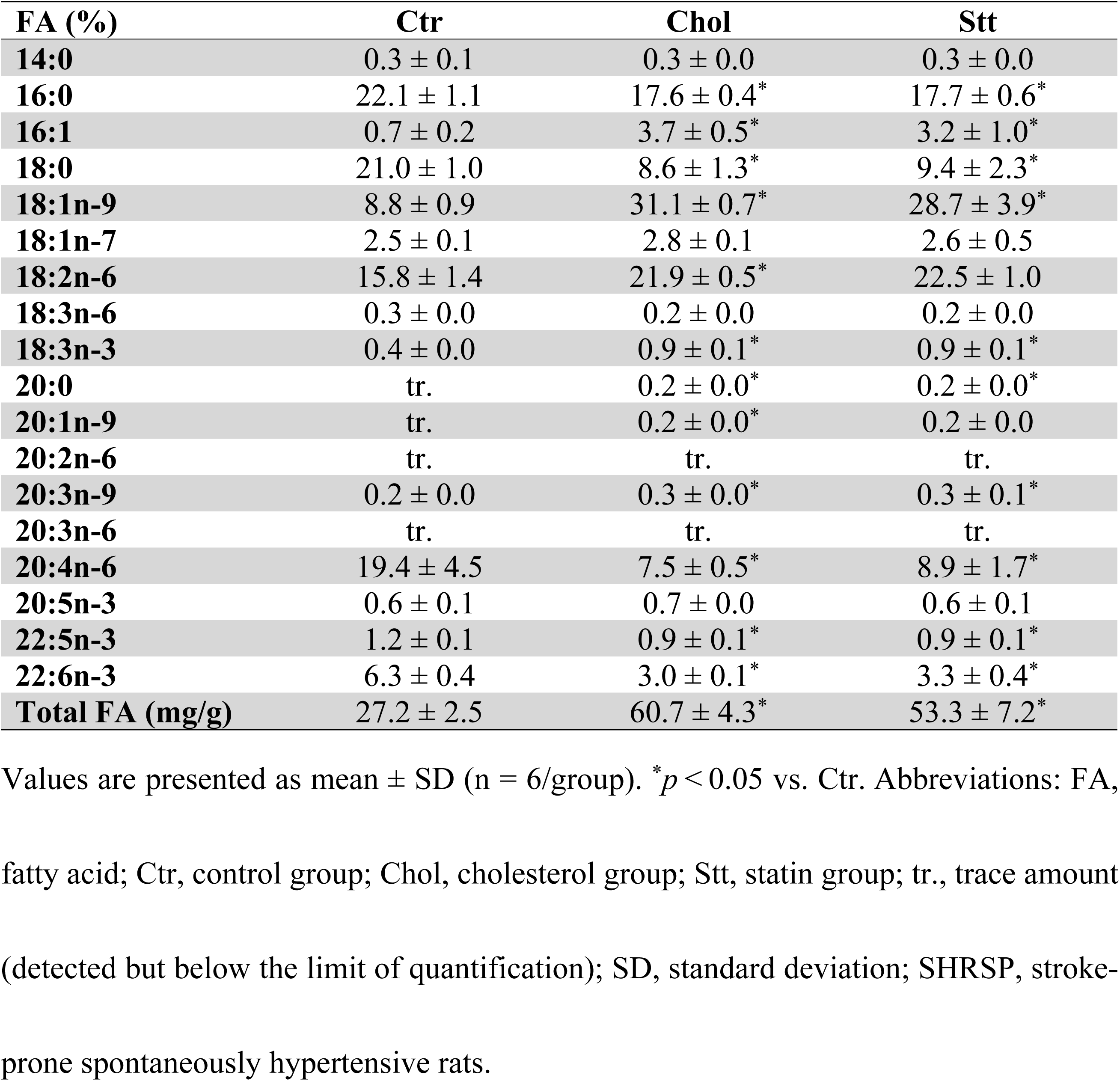
FA compositions and levels in the liver of SHRSP rats.

**Table 6.**
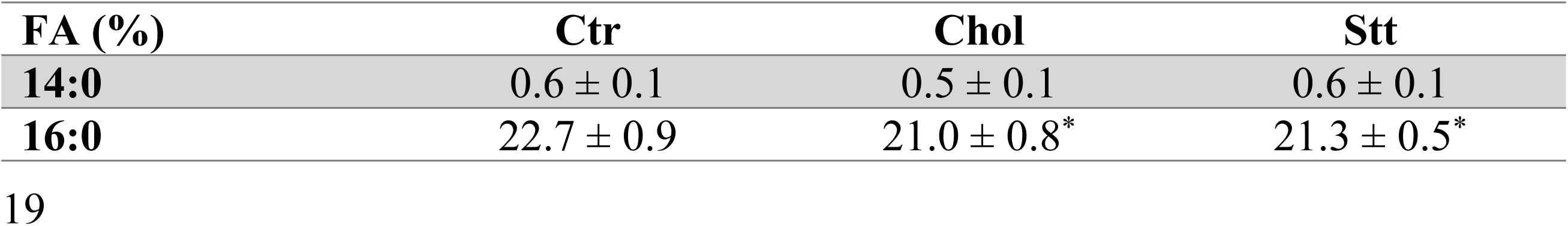
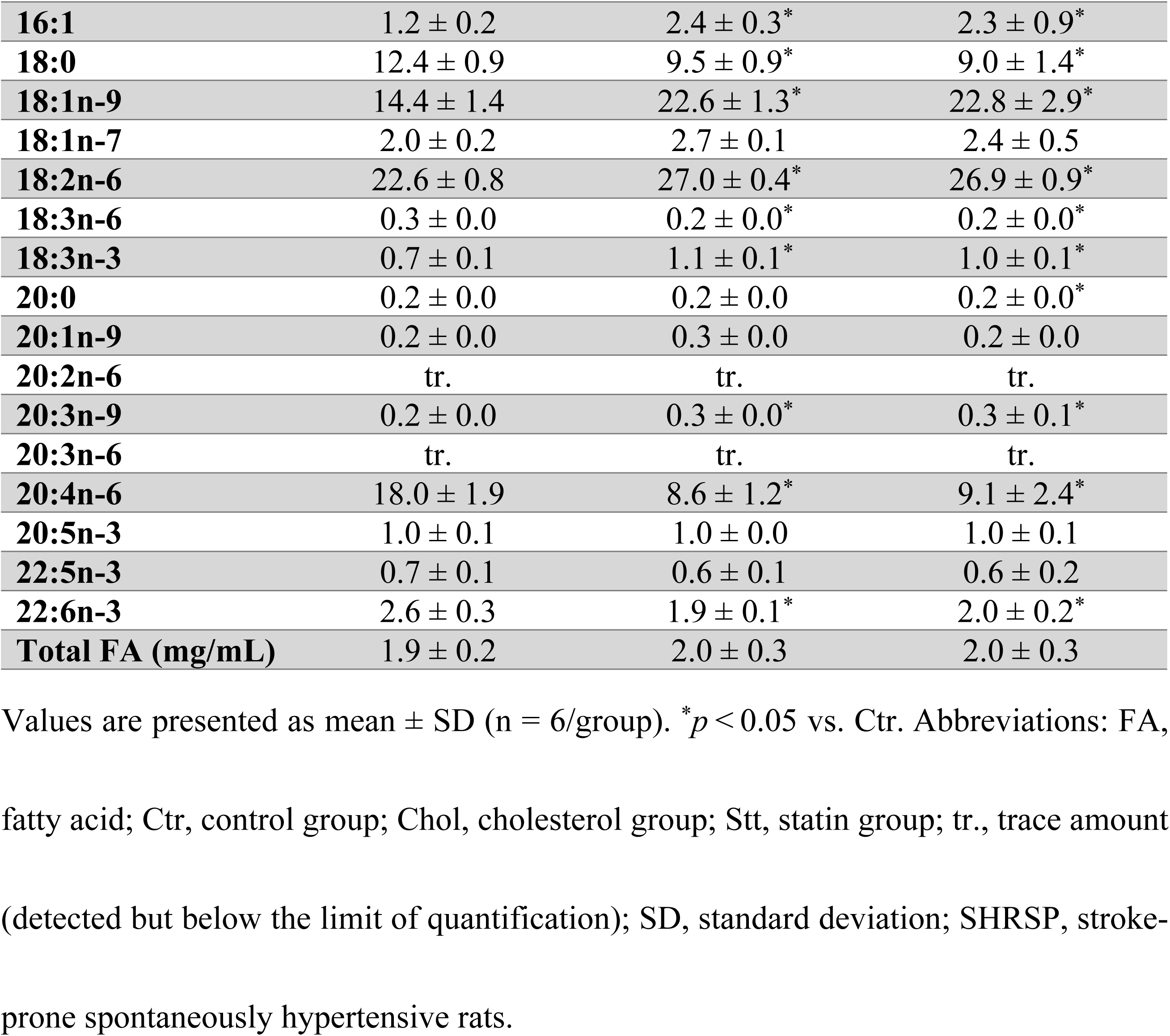
FA compositions and levels in serum of SHRSP rats.

| FA (%) | Ctr | Chol | Stt |
| --- | --- | --- | --- |
| **14:0** | 0.6 ± 0.1 | 0.5 ± 0.1 | 0.6 ± 0.1 |
| **16:0** | 22.7 ± 0.9 | 21.0 ± 0.8* | 21.3 ± 0.5* |
| **16:1** | 1.2 ± 0.2 | 2.4 ± 0.3* | 2.3 ± 0.9* |
| **18:0** | 12.4 ± 0.9 | 9.5 ± 0.9* | 9.0 ± 1.4* |
| **18:1n-9** | 14.4 ± 1.4 | 22.6 ± 1.3* | 22.8 ± 2.9* |
| **18:1n-7** | 2.0 ± 0.2 | 2.7 ± 0.1 | 2.4 ± 0.5 |
| **18:2n-6** | 22.6 ± 0.8 | 27.0 ± 0.4* | 26.9 ± 0.9* |
| **18:3n-6** | 0.3 ± 0.0 | 0.2 ± 0.0* | 0.2 ± 0.0* |
| **18:3n-3** | 0.7 ± 0.1 | 1.1 ± 0.1* | 1.0 ± 0.1* |
| **20:0** | 0.2 ± 0.0 | 0.2 ± 0.0 | 0.2 ± 0.0* |
| **20:1n-9** | 0.2 ± 0.0 | 0.3 ± 0.0 | 0.2 ± 0.0 |
| **20:2n-6** | tr. | tr. | tr. |
| **20:3n-9** | 0.2 ± 0.0 | 0.3 ± 0.0* | 0.3 ± 0.1* |
| **20:3n-6** | tr. | tr. | tr. |
| **20:4n-6** | 18.0 ± 1.9 | 8.6 ± 1.2* | 9.1 ± 2.4* |
| **20:5n-3** | 1.0 ± 0.1 | 1.0 ± 0.0 | 1.0 ± 0.1 |
| **22:5n-3** | 0.7 ± 0.1 | 0.6 ± 0.1 | 0.6 ± 0.2 |
| **22:6n-3** | 2.6 ± 0.3 | 1.9 ± 0.1* | 2.0 ± 0.2* |
| **Total FA (mg/mL)** | 1.9 ± 0.2 | 2.0 ± 0.3 | 2.0 ± 0.3 |

Values are presented as mean ± SD (n = 6/group). \**p* < 0.05 vs. Ctr. Abbreviations: FA, fatty acid; Ctr, control group; Chol, cholesterol group; Stt, statin group; tr., trace amount (detected but below the limit of quantification); SD, standard deviation; SHRSP, stroke- prone spontaneously hypertensive rats.

**Table 7.** FA composition and levels of phospholipids in the kidney of SHRSP rats.

| FA (%) | Ctr | Chol | Stt |
| --- | --- | --- | --- |
| 14:0 | 0.3 ± 0.0 | 0.2 ± 0.0 | 0.2 ± 0.0 |
| 16:0 DMA | 2.3 ± 0.1 | 2.2 ± 0.1 | 2.3 ± 0.1 |
| 16:0 | 26.1 ± 1.7 | 24.7 ± 0.5* | 24.9 ± 0.3* |
| 16:1 | 0.4 ± 0.1 | 0.6 ± 0.2* | 0.4 ± 0.0<sup>#</sup> |
| 18:0 DMA | 1.5 ± 0.1 | 1.3 ± 0.0* | 1.4 ± 0.1* |
| 18:0 | 19.8 ± 0.3 | 19.1 ± 0.7 | 19.1 ± 0.4 |
| 18:1n-9 | 7.9 ± 0.3 | 8.7 ± 0.5* | 8.7 ± 0.4* |
| 18:1n-7 | 1.5 ± 0.0 | 1.7 ± 0.1* | 1.7 ± 0.0* |
| 18:2n-6 | 7.5 ± 0.3 | 10.9 ± 0.6* | 10.6 ± 0.3* |
| 18:3n-6 | tr. | tr. | tr. |
| 18:3n-3 | 0.3 ± 0.0 | 0.3 ± 0.0 | 0.3 ± 0.1 |
| 20:0 | 0.2 ± 0.0 | 0.2 ± 0.0 | 0.2 ± 0.0 |
| 20:1n-9 | tr. | tr. | tr. |
| 20:2n-6 | tr. | tr. | tr. |
| 20:3n-9 | tr. | 0.2 ± 0.0* | 0.2 ± 0.0* |
| 20:3n-6 | tr. | tr. | tr. |
| 20:4n-6 | 28.7 ± 0.2 | 26.0 ± 0.4* | 26.5 ± 0.4* |
| 20:5n-3 | 0.4 ± 0.0 | 0.6 ± 0.2* | 0.5 ± 0.0 |
| 22:0 | 0.5 ± 0.0 | 0.4 ± 0.0 | 0.5 ± 0.1 |
| 22:5n-3 | 0.3 ± 0.0 | 0.3 ± 0.0 | 0.3 ± 0.0 |
| 22:6n-3 | 2.0 ± 0.2 | 2.2 ± 0.6 | 1.8 ± 0.1 |
| Total FA (mg/g) | 16.2 ± 0.7 | 16.3 ± 1.6 | 15.2 ± 1.0 |

Values are presented as mean ± SD (n = 5-6/group). \**p* < 0.05 vs. Ctr; <sup>#</sup>*p* < 0.05 vs.
Chol. Abbreviations: FA, fatty acid; Ctr, control group; Chol, cholesterol group; Stt,
statin group; DMA, dimethyl acetal; tr., trace amount (detected but below the limit of
quantification); SD, standard deviation; SHRSP, stroke-prone spontaneously
hypertensive rats.

### Biochemistry

Fig 4A shows that hepatic PGE₂ levels were decreased in both the Chol and Stt groups, and a similar, though nonsignificant, decreasing trend was observed in the kidney (Fig 4B).

**Fig 4.**
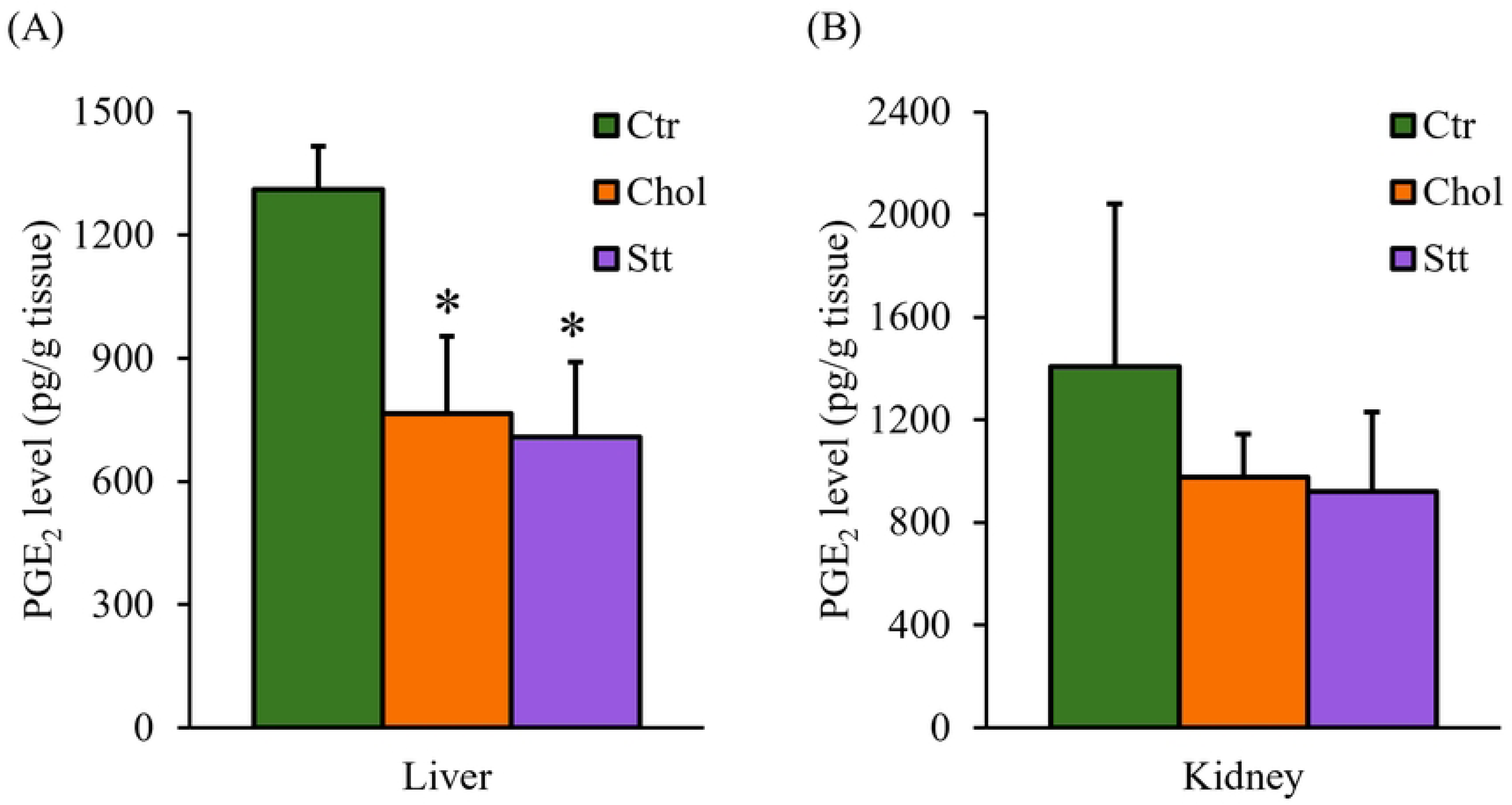
PGE_2_ levels in the liver and kidney of SHRSP rats. (A) Hepatic and (B) renal PGE_2_ levels. Values are presented as mean ± SD (n = 6/group). \**p* < 0.05 vs. Ctr. Abbreviations: Ctr, control group; Chol, cholesterol group; Stt, statin group; SD, standard deviation; PGE_2_, prostaglandin E_2_; SHRSP, stroke-prone spontaneously hypertensive rats.

Fig 5A shows that hepatic SOD activity was lower in both the Chol and Stt groups than in the Ctr group. Serum SOD activity was increased in the Chol group compared with the Ctr group, whereas there was no significant difference between the Ctr and Stt groups (Fig 5B). Renal SOD activity was not significantly affected by either diet (Fig 5C).

**Fig 5.**
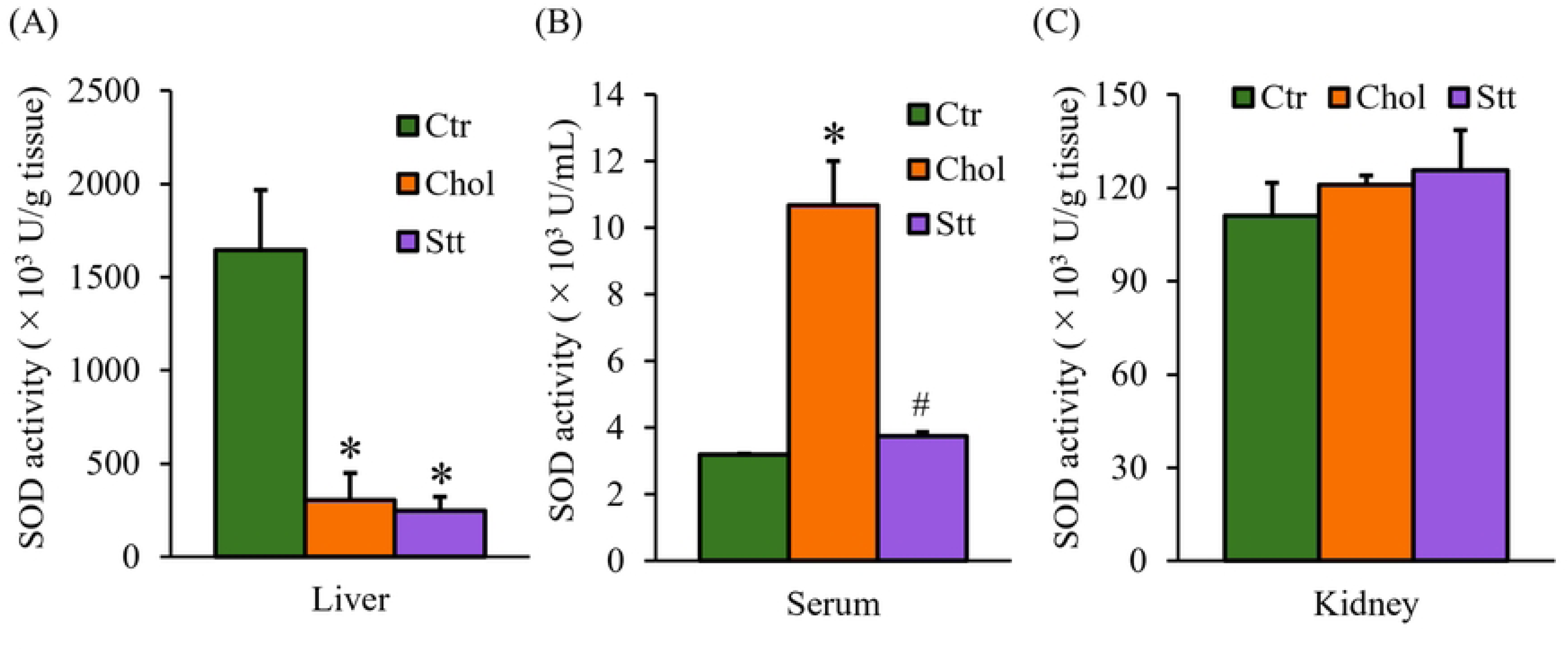
SOD activities in the liver, serum, and kidney of SHRSP rats. (A) Hepatic, (B) serum, and (C) renal SOD activities. Values are presented as mean ± SD (n = 2/group). One unit (U) of SOD is defined as the amount of the enzyme that inhibits the reduction reaction of WST-1 with superoxide anion by 50%. \**p* < 0.05 vs. Ctr; ^#^*p* < 0.05 vs. Chol. Abbreviations: Ctr, control group; Chol, cholesterol group; Stt, statin group; SD, standard deviation; SOD, Superoxide dismutase; SHRSP, stroke-prone spontaneously hypertensive rats.

### mRNA expression

The hepatic mRNA expression levels are exhibited in Figure 6. The analysis revealed that the genes implicated in sterol transport (*Abcg5, Abcg8*, and *Npc1l1*) exhibited a more than twofold increase in the Chol group compared with the Ctr group. Conversely, these genes demonstrated a decrease in expression levels in the Stt group (Fig. 6A). The expression of *Hmgcr* was found to be significantly reduced in both the Chol and Stt groups; however, a more pronounced decrease was observed in the Stt group. The low-density lipoprotein receptor (*Ldlr*) exhibited no substantial variance between the Ctr and Chol groups; however, a considerable decrease was evident in the Stt group in comparison to the Chol group.

**Fig 6.**
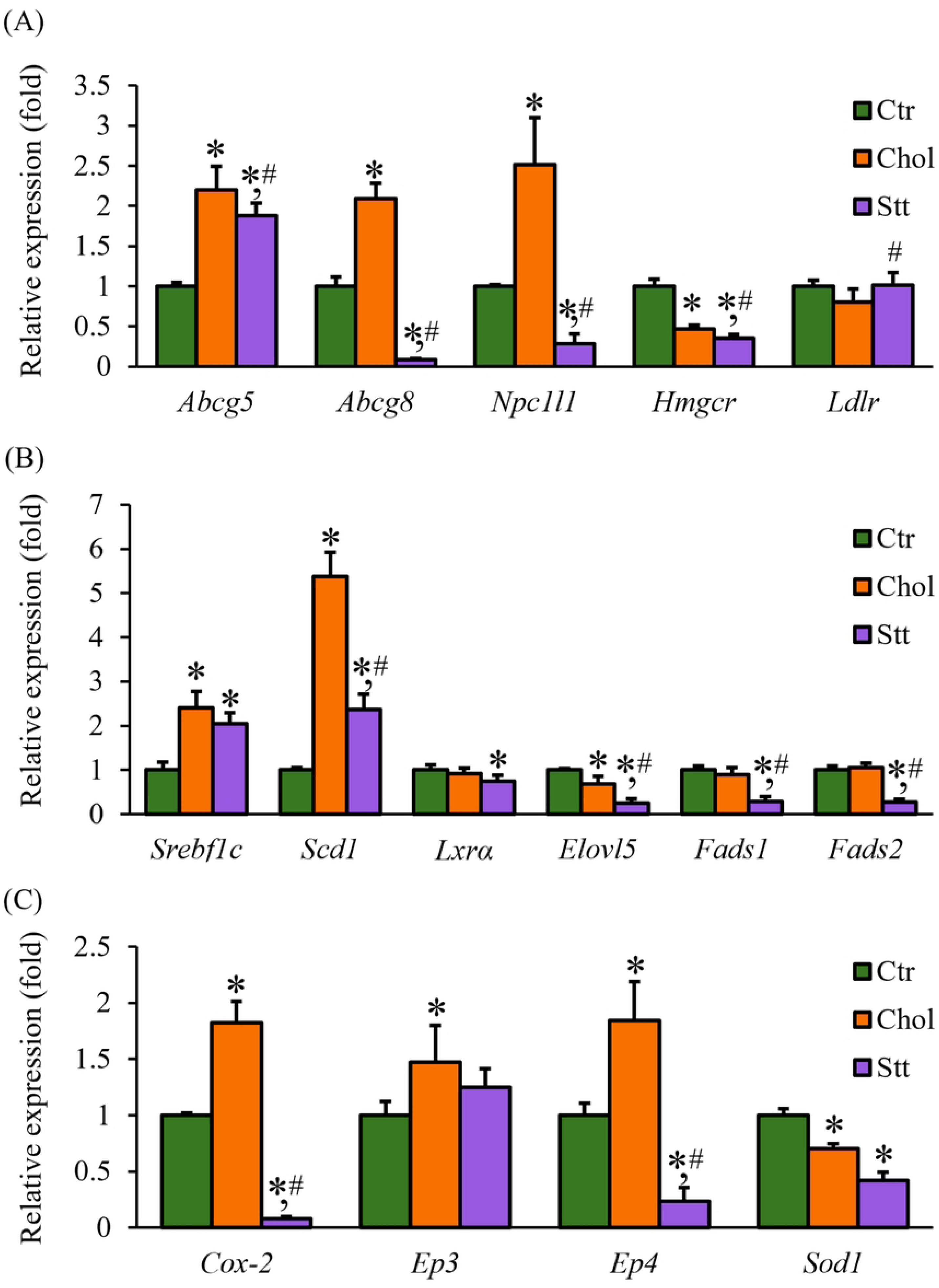
Hepatic mRNA expression of lipid metabolism–related genes in SHRSP rats. Hepatic mRNA expression related to (A) sterol metabolism, (B) FA metabolism, and (C) others. Values are presented as mean ± SD (n = 6/group). \**p* < 0.05 vs. Ctr; ^#^*p* < 0.05 vs. Chol. Abbreviations: Ctr, control group; Chol, cholesterol group; Stt, statin group; SD, standard deviation; *Abcg5*, ATP-binding cassette sub-family G member 5; *Abcg8*, ATP-binding cassette sub-family G member 8; *Npc1l1*, Niemann-Pick C1-like 1; *Hmgcr*, 3-hydroxy-3-methyl-glutaryl-coenzyme A (HMG-CoA) reductase; *Ldlr*, Low density lipoprotein receptor, *Srebf1c*, Sterol regulatory element-binding transcription factor 1c; *Scd1*, Stearoyl-CoA desaturase 1; *Lxrα*, Liver X receptor α; *Elovl5*, Elongation of very long chain fatty acids 5; *Fads1*, Fatty acid desaturase 1; *Fads2*, Fatty acid desaturase 2; *Cox-2*, Cyclooxygenase-2; *Ep3*, Prostaglandin E_2_ receptor 3, *Ep4*, Prostaglandin E_2_ receptor 4; *Sod1*, Superoxide dismutase 1; SHRSP, stroke-prone spontaneously hypertensive rats.

Among the genes implicated in FA metabolism, sterol regulatory-element binding factor 1c (*Srebf1c*) and stearoyl-CoA desaturase 1 (*Scd1*) exhibited increased expression levels in the Chol group (Fig. 6B). In the Stt group, *Scd1* expression levels were lower than those observed in the Chol group; however, they remained higher than those seen in the Ctr group. Furthermore, the expression of liver X receptor α (*Lxrα*) did not exhibit a significant difference between the Ctr and Chol groups; however, it demonstrated a significant decrease in the Stt group compared with the Ctr group. The expression of elongation of very long chain fatty acids 5 *(Elovl5*) was found to be decreased in the Chol group, whereas the expression levels of fatty acid desaturase 1 (*Fads1*) and *Fads2* remained constant. Conversely, *Fads1, Fads2*, and *Elovl5* exhibited a marked decrease in the Stt group.

With respect to the AA–PGE_2_ pathway, the expression levels of cyclooxygenase 2 (*Cox-2*), prostaglandin E2 receptor 3 (*Ep3*), and Ep4 were increased in the Chol group compared with the Ctr group by approximately 1.8-, 1.4-, and 1.8-fold, respectively. Conversely, *Cox-2* and *Ep4* were decreased in the Stt group (Fig. 6C). The expression of superoxide dismutase 1 (*Sod1*) was found to be reduced in both diet groups.

### Histological analysis

The evaluation of kidney sections that had been stained with hematoxylin and eosin is illustrated in Figure 7 and detailed in Table 8. Slight infiltration of inflammatory cells in glomeruli (+) was observed in three of six rats in the Ctr group, whereas only very slight infiltration (±) was observed in the Chol and Stt groups. The presence of inflammatory cells in the interstitium (+) was observed in one rat in the Ctr group. However, basophilia in the tubule and cortex (+) was observed in one or two rats in each group, and the grades of granular cast and hyaline cast in the tubule did not exceed very slight (±) in any group.

**Fig 7.**
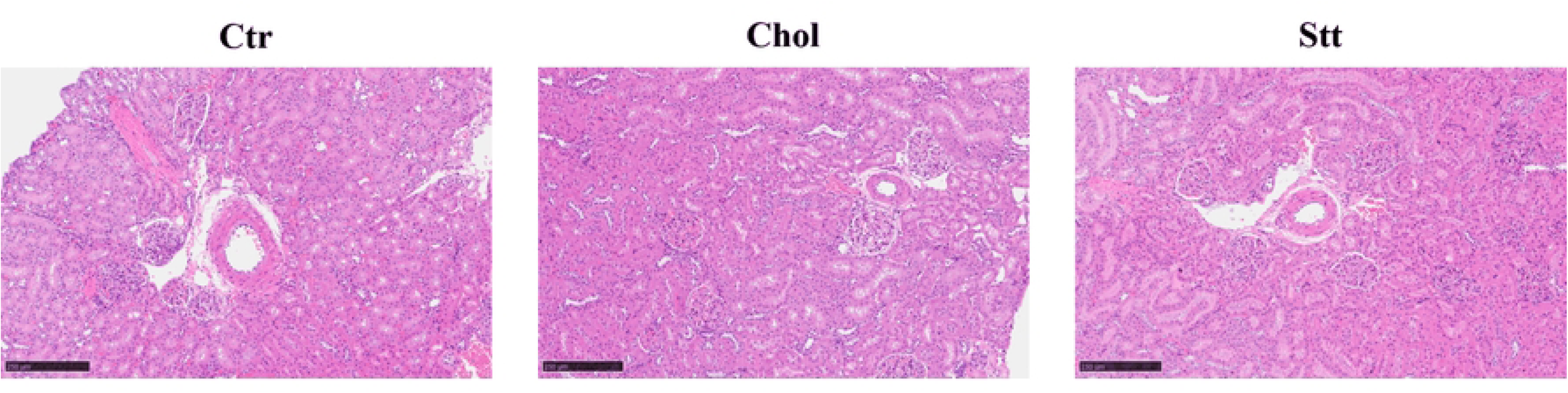
Histological analysis of the kidney samples from SHRSP. Tissue sections were stained with hematoxylin and eosin. Scale bar = 250 µm. Abbreviations: Ctr, control group; Chol, cholesterol group; Stt, statin group; SHRSP, stroke-prone spontaneously hypertensive rats.

**Table 8.** Summary of the histological analysis of the kidney from SHRSP.

|  | **Ctr** | **Chol** | **Stt** |
| --- | --- | --- | --- |
| **Infiltrate, inflammatory cell, glomeruli** | (±)×3, (+)×3 | (±)×6 | (±)×6 |
| **Basophilia, tubule, cortex** | (±)×4, (+)×2 | (±)×5, (+)×1 | (±)×4, (+)×1 |
| **Cast, hyaline, tubule** |  | (±)×2 | (±)×1 |
| **Cast, granular, tubule** | (±)×1 |  | (±)×2 |
| **Infiltrate, inflammatory cell, interstitium** | (±)×5, (+)×1 | (±)×6 | (±)×6 |

Values indicate the number of rats in each histological grade. Histological changes were evaluated using a five-point scale (–, ±, +, ‡, †), and represents the number scored more than (±). Abbreviations: Ctr, control group; Chol, cholesterol group; Stt, statin group; SHRSP, stroke-prone spontaneously hypertensive rats.

## Discussion

Chol is an essential biological molecule that plays a critical role in the construction of cell membranes and the synthesis of steroid hormones and bile acids [1]. A high dietary Chol intake has been demonstrated to be associated with an increased risk of cardiovascular diseases, including atherosclerosis [2, 24]. Conversely, low serum Chol levels have been associated with an elevated risk of hemorrhagic stroke in humans [3]. SHRSP rats demonstrate persistently low serum Chol levels, largely due to reduced hepatic Chol synthesis when compared with normotensive WKY rats [25]. This hypocholesterolemic state has been proposed as a factor contributing to their tendency to develop hemorrhagic stroke. A prior study has indicated that SHRSP rats exhibit a lifespan benefit when fed a diet augmented with 1% w/w Chol [26]. The present study sought to elucidate the potential role of dietary Chol in promoting lifespan extension in SHRSP rats through biochemical and histological analyses.

Previous research has indicated that stroke development in SHRSP rats is accompanied by a decline in body weight and food intake [27]. In the present study, no such changes were observed among the three diet groups after 12 weeks of feeding (Fig. 1). Conversely, systolic blood pressure (SBP) exhibited a marked decrease in the Chol and Stt groups compared with the Ctr group following 12 weeks (Fig. 2). These findings suggest that dietary Chol may lower SBP in SHRSP rats and thereby diminish the risk of hemorrhagic events.

The lifespan of SHRSP rats is contingent upon the consumption of various fats and oils [6–8,27]. A multitude of studies have indicated that oils with high concentrations of PS have the potential to reduce their longevity [10]. PS are plant-derived sterols with structures similar to Chol but with differences in the length and degree of saturation of their side chains. The precise mechanisms underlying PS-induced pathology in SHRSP rats remain to be elucidated. However, PS is incorporated into cell membranes, a process comparable to that of Chol, and has the capacity to modify membrane properties [10]. Intake of rapeseed oil, which is rich in PS, has also been shown to increase blood pressure in SHRSP rats. This increase in blood pressure was accompanied by suppression of Na^+^/K^+^-ATPase activity. This suggests a link between membrane sterol composition and blood pressure regulation [28]. Chol and PS have been observed to share intestinal absorption pathways via NPC1L1. Consequently, an increase in dietary Chol intake has the potential to suppress PS absorption and reduce tissue PS levels [12, 13]. In accordance with these findings, the present study demonstrated that feeding a Chol-enriched diet led to a decrease in PS levels within the kidney (Fig. 3C). This reduction in renal PS content may contribute to improved membrane properties and blood pressure regulation in SHRSP rats (Fig. 2), although this remains speculative. The regulation of sterol transport is achieved through a dual mechanism: NPC1L1-mediated uptake and ABCG5/8-mediated biliary excretion [13]. The expression levels of *Npc1l1* and *Abcg5/8* mRNA were elevated in the liver following 12 weeks of Chol feeding (Fig. 6A), indicating an augmented sterol trafficking process. However, SHRSP rats harbor a mutation in *Abcg5*, which impairs sterol excretion and predisposes them to hepatic sterol accumulation. Consequently, an augmented uptake in the context of defective excretion may have promoted hepatic sterol accumulation.

It has been reported that Sito and Camp act as LXRα ligands and activate the receptor in certain cell lines [29]. While the present study did not directly assess LXRα activity, the increase in total PS content observed in the liver (Fig. 3A) may be indicative of agonist activity at LXRα, which contributed to the rise in FA content. Furthermore, PS is oxidized by oxidative stress and other factors to form oxidized PS (OPS) [30]. Recent studies have revealed that the ingestion of certain forms of OPS can suppress the sterol synthesis pathway and enhance the FA synthesis pathway, independently of micelle formation and Chol competition [31]. Although OPS was not directly measured in this study, it is possible that a fraction of PS was oxidized to OPS in vivo under oxidative stress, thereby enhancing FA synthesis via a distinct pathway.

It has been established that disparities in the side-chain configuration of PSs exert a significant influence on their micellar solubility [32] and have been posited to modulate micellar affinity [33]. These physicochemical distinctions may underlie the divergent effects exhibited by Sito and Camp. In healthy individuals, Sito has been shown to have lower intestinal absorption rates (4%–8%) compared to Camp (9%–18%) and approximately 2.6 times higher hepatic clearance, suggesting that Camp is less readily excreted than Sito [34]. A comparable disparity in hepatic clearance may exist in the SHRSP rats employed in this study, indicating the possibility of liver-specific Camp accumulation (see Fig. 3A-C). However, despite the increase in hepatic Camp, its levels in serum and kidney were lower in both Chol and Stt groups. This finding suggests that Camp is retained within the liver rather than released into circulation in SHRSP rats, although the underlying mechanism remains unclear. In recent years, it has been revealed that in Abcg5/8 knockout female mice, Sito exhibits reproductive toxicity in the ovaries due to epigenetic modifications, whereas no such significant effects are observed in Camp and Stig [35]. However, many aspects of the functional differences in vivo associated with different PS molecular species remain unclear. Further investigation is necessary to understand the impact of the PS accumulation differences observed in this experiment on stroke onset.

In terms of FA composition, the AA ratio was decreased in phospholipids from serum, liver, and kidney in the Chol and Stt groups (Table 5-7). AA is a precursor of eicosanoids, and some of them, such as PGE_2_, are lipid mediators that regulate inflammatory response [21]. Reduced AA availability is known to decrease the production of pro-inflammatory mediators, thereby suppressing inflammation and tissue injury in various disease models and clinical settings [36]. Consistent with this concept, hepatic and renal PGE₂ levels showed a decreasing tendency accompanied by a reduction in AA following ingestion of a Chol-rich diet in the present study (Table 5 and Fig 4). Previous studies have demonstrated that docosahexaenoic acid (22:6n-3; DHA) administration suppresses the development of hypertension in SHRSP rats by lowering plasma AA levels [37]. Although the mechanism by which Chol reduces AA levels might differ from that of DHA, the observed decrease in AA induced by Chol intake may similarly contribute to attenuation of SHRSP pathogenesis.

The induction of FA synthetic enzymes is subject to regulation via the SREBP-1c pathway, which is located downstream of LXRα [19]. Ligands for LXRα include oxidized Chol (oxiChol), which is produced through CYP-mediated oxidation of Chol [38]. In this experiment, *Lxrα* mRNA expression did not differ significantly between the Chol and Ctr groups and was slightly lower in the Stt group (Fig. 6B). Conversely, *Srebf1c* (encoding SREBP-1c) expression exhibited an increase in both the Chol and Stt groups. Although oxiChol levels were not directly measured, increased production of oxiChol following Chol intake may have activated LXRα at the functional level, thereby enhancing SREBP-1c expression without altering *Lxrα* mRNA abundance.

AA biosynthesis involves FADS2, ELOVL5, and FADS1; *Elovl5* mRNA expression was decreased in both the Chol and Stt groups (Fig. 6B). This finding appears inconsistent with the upregulation of *Srebf1c*, which is known to induce *Elovl5* transcription [39]. *Elovl5* expression has been documented to be repressed in murine models of fatty liver disease under specific conditions [40]. Consequently, the observed decline may be linked to hepatic lipid accumulation, as evidenced by increased liver weight and hepatic total FA and Chol content. However, the precise regulatory mechanisms underlying this phenomenon remain to be elucidated. However, the observed decrease in *Elovl5* is likely to have contributed to the decline in AA, which in turn resulted in reduced PGE_2_ levels.

PGE₂ exerts its biological effects through four receptors, designated EP1-4 [41]. Of these, EP3 is predominantly linked to pro-inflammatory signaling [42], while EP4 has been implicated in anti-inflammatory and immunosuppressive responses, including protection against cardiovascular inflammation [43,44]. In the present study, the expression levels of *Cox-2* and both *Ep3* and *Ep4* receptors were found to be increased; however, the relative increase in *Ep4* receptor was greater than that of *Ep3* receptor (Fig. 6C). In conjunction with the observed decrease in PGE₂ production (see Figure 4), these findings indicate that anti-inflammatory signaling pathways were predominantly active in the liver following the ingestion of Chol. If analogous alterations occur in other organs, then it is possible that Chol consumption may have a systemic anti-inflammatory effect on SHRSP rats.

Despite the findings of prior studies indicating that diets abundant in oleic acid result in a reduction of lifespan in SHRSP rats [6,7], oleic acid levels were elevated in both the liver and serum in the present study (Tables 5 and 6). This study was conducted under conditions reported to extend the lifespan of SHRSP rats. These findings imply that alterations in FA metabolic pathways, rather than the intake of a specific FA per se, may be involved in regulating disease progression.

During the metabolism of Chol and FAs, oxidative reactions catalyzed by cytochrome P450 and COX generate reactive oxygen species (ROS) [45]. In the present study, Chol intake reduced hepatic *Sod1* mRNA expression and SOD activity (Fig 5A and 6C). This indicates a decline in local antioxidant capacity, possibly due to increased oxidative stress associated with fatty liver development and Chol accumulation. Taken together, the present findings suggest that Chol intake may induce hepatic oxidative stress and functional impairment, even as it exerts protective effects against stroke development.

Rapeseed oil intake has been reported to shorten lifespan and reduce erythrocyte SOD activity in SHRSP rats [46,47]. In contrast, in the present study, serum SOD activity was significantly increased in the Chol group after 12 weeks of feeding (Fig 5B). This elevation in antioxidant capacity may have enhanced resistance to oxidative stress and attenuated tissue damage. Such an increase in circulating antioxidant defense may therefore be associated with improved pathology and lifespan extension in Chol-fed SHRSP rats.

In SHRSP rats, hypertension caused by renal impairment is considered a contributor to fatal stroke [48]. Histological examination of renal tissue revealed glomerular deformation in the Ctr group, whereas this change was relatively mild in the Chol and Stt groups (Fig 7). Infiltration of inflammatory cells was slight (+) in three cases in the Ctr group and very slight (±) in all cases in the Chol and Stt groups (Table 8). Periglomerular cells play a key role in blood pressure regulation by regulating renin secretion [49]. The renal lesions observed in the Ctr group likely reflect pathological changes associated with progressive hypertension, which appear to have been suppressed by Chol intake. The protective effect of Chol may involve reduced PS accumulation and inhibition of the AA–PGE₂ pathway. These changes may contribute to the preservation of renal function and stabilization of blood pressure regulation, thereby contributing to improved pathology and lifespan extension [27,50].

Stt inhibits HMGCR and lowers circulating Chol by upregulating LDLR [51]. In the present study, a lifespan-extending dose of Stt was added to a 1% high-Chol diet to examine the effects of combined administration. In the Stt group, *Hmgcr* expression was decreased, whereas *Ldlr* expression was increased compared with the Chol group, indicating that Stt modified hepatic Chol metabolism even under conditions of high dietary Chol intake (Fig 6A). Although sterol profiles (Fig 3), serum SOD activity (Fig 5), and FA-related gene expression (Fig 6B and C) differed between the Stt and Chol groups, no significant differences were observed in renal PS content (Fig 3C), AA ratio (Table 7), PGE₂ levels (Fig 4), histological changes (Fig 7 and Table 8), or blood pressure suppression (Fig 2). These findings suggest that, under a high-Chol diet (1% w/w), a standard dose of Stt might not further enhance the lifespan-extending effects of Chol intake.

Both the Stt and Chol groups demonstrated substantial improvements in pathology compared with the Ctr group. These results indicate that dietary Chol intake plays a predominant role in pathological improvement, even when endogenous Chol synthesis is pharmacologically suppressed. This finding suggests that dietary and endogenous Chol may have distinct contributions to disease progression in SHRSP rats.

While dietary Chol is generally regarded as a risk factor for cardiovascular disease [24], the present study indicates that Chol may also contribute to physiological homeostasis beyond its pathological effects. A key strength of this study is the experimental design, which involved supplementing a standard diet with exogenous Chol without altering dietary FA composition. This approach allowed for the specific assessment of dietary Chol effects independent of FA intake, a methodology rarely employed in previous research. Additionally, the use of SHRSP rats, which possess a genetic predisposition affecting sterol metabolism, facilitated a detailed examination of the relationships among Chol intake, lipid metabolism, renal pathology, and stroke-related outcomes. These findings provide mechanistic insights into how dietary Chol may reduce the risk of hemorrhagic stroke in genetically susceptible populations.

Several limitations of this study should be acknowledged. First, the dietary Chol content used in this experiment (1% w/w) exceeds typical human intake levels and was administered under conditions without additional dietary FAs. Therefore, while this design is advantageous for mechanistic analysis, the findings cannot be directly extrapolated to ordinary human diets, in which Chol and FAs are consumed simultaneously. Second, SHRSP rats harbor an *Abcg5* mutation and exhibit distinct sterol metabolism, which may further limit direct application of these results to humans or other animal models. Third, the present analysis examined only a single statin dose under high-Chol dietary conditions; thus, dose-dependent interactions between dietary Chol and statin treatment cannot be excluded. Finally, although alterations in sterol profiles, FA metabolism, and antioxidant capacity were observed, key intermediates and signaling molecules were not directly measured, and the precise molecular mechanisms remain to be fully elucidated. Despite these limitations, the present study clarifies how dietary Chol modulates lipid metabolism, renal pathology, and stroke-related outcomes under conditions of genetic susceptibility.

To address these limitations, future research should include dose-response studies to clarify the effects of varying levels of dietary Chol and statin intake on lipid metabolism and blood pressure regulation. Additional studies using animal models without genetic predispositions would help confirm these findings and broaden their applicability. Furthermore, mechanistic assays focusing on specific signaling pathways affected by dietary Chol could elucidate the underlying processes governing lipid metabolism and renal pathology. Longitudinal studies investigating chronic dietary Chol intake and its effects on cardiovascular health over extended periods would also offer important perspectives.

## Conclusions

The present study demonstrated that the Chol diet limits blood pressure elevation in SHRSP rats and suppresses PS accumulation, the AA–PGE₂ pathway, and renal histopathological changes. These effects appear to be mediated through specific mechanistic pathways involving sterol transport, AA metabolism, and antioxidant defense. Chol intake appears to reduce systolic blood pressure by altering sterol transport, decreasing AA-derived pro-inflammatory mediators, and enhancing antioxidant capacity. Suppression of the AA–PGE₂ pathway indicates a shift toward anti-inflammatory responses. Additionally, changes in sterol composition, which reduce membrane fragility, may contribute to blood pressure regulation and renal protection. By isolating the effects of exogenous Chol from those of dietary FAs, this study provides novel insights into the physiological roles of Chol intake under stroke-prone conditions. Although extrapolating these findings to humans requires caution, they suggest that dietary Chol intake may modulate disease-related pathological features in susceptible individuals.

## Author contributions

Y.N. Formal analysis, Investigation, Writing–original draft.

K.T. Conceptualization, Methodology, Formal analysis, Investigation, Resources, Writing–review & editing draft, Visualization, Supervision, and Funding acquisition.

Y.S. Investigation and supervision for tissue fixing and histological analysis.

T.M. Supervising investigators & analysis.

N.O. Supervising histological analysis & draft.

## Acknowledgements

The authors thank S.N., who helps to maintaine the animal experiments.

## Supporting information

**S1 Table. Data of weight, food intake, organ weight, and systolic blood pressure**

**S2 Table. Data of FA analyses**

**S3 Table. Data of sterol analyses**

**S4 Table. Data of other biochemical analyses**

**S5 Table. Data of real-time RT-PCR**

